# dcSBM: A federated constrained source-based morphometry approach for multivariate brain structure mapping

**DOI:** 10.1101/2022.12.29.522266

**Authors:** Debbrata K. Saha, Rogers F. Silva, Bradley T. Baker, Rekha Saha, Vince D. Calhoun

## Abstract

The examination of multivariate brain morphometry patterns has gained attention in recent years, especially for their powerful exploratory capabilities in the study of differences between patients and controls. Among many existing methods and tools for analysis of brain anatomy based on structural magnetic resonance imaging (sMRI) data, data-driven source based morphometry (SBM) focuses on the exploratory detection of such patterns. Constrained source-based morphometry (constrained SBM) is a widely used semi-blind extension of SBM that enables extracting maximally independent reference-alike sources using the constrained independent component analysis (ICA) approach. In order to operate, constrained SBM needs the data to be locally accessible. However, there exist many reasons (e.g., the concerns of revealing identifiable rare disease information, or violating strict IRB policies) that may preclude access to data from different sites. In this scenario, constrained SBM fails to leverage the benefits of decentralized data. To mitigate this problem, we present a novel approach: decentralized constrained source-based morphometry (dcSBM). In dcSBM, the original data never leaves the local site. Each site operates constrained ICA on their private local data while using a common distributed computation platform. Then, an aggregator/master node aggregates the results estimated from each local site and applies statistical analysis to find out the significant sources. In our approach, we first use UK Biobank sMRI data to investigate the reliability of our dcSBM algorithm. Finally, we utilize two additional multi-site patient datasets to validate our model by comparing the resulting group difference estimates from both centralized and decentralized constrained SBM.

## 1. Introduction

Structural magnetic resonance imaging(sMRI) is a widely used neuroimaging technique which can be used to assess brain morphometry. A traditional approach to extract brain structure changes is to segment the brain into regions of interest (ROI) and finally compute the difference between groups at each ROI. However, this approach makes the assumption that the brain can be divided anatomically from the MRI data. There is another approach called Voxel based morphometry (VBM) which is used to estimate the voxel-wise differences [1]. As a massively univariate approach, VBM reports voxel-wise changes across the brain in the form of statistical maps. However, its univariate nature means it can not extract information about the relationship among the voxels. Also, VBM estimates are limited to the specific predictive effect defined by the user/design. In order to capture the inter-voxel information, a multivariate approach would be preferred.

Source based morphometry (SBM) is a multivariate data-driven approache introduced to automatically detect whole brain structure by using the information over voxels [2]. The goal of SBM is to estimate maximally spatially independent sources (whole-brain spatial maps) that covary among subjects, and to compute potential individual subject or group differences based on the degree of expression of these sources. SBM utilizes the combined techniques of VBM and independent component analysis (ICA) to extract the spatially independent sources, and uses statistical measures to find out the source of interest. Recent studies report that SBM can preserve the spatial correlation between different brain regions [3, 4]. SBM can be used to analize different type of disorders [5] such as Parkinson’s [6], bipolar and borderline personality disorders [7], schizophrenia and bipolar disorder [8], as well as brain abnormalities in children with autism [9, 10].

Combining both ROI and data driven methodologies, another mulitvariate data-driven approach was proposed, namely, constrained source-based morphometry (cSBM) [11]. cSBM leverages the full utility of SBM with the ROI information, and also provides automatic correspondence between datasets. In constrained SBM, a prior ROI or a whole-brain reference is used as a constraint in the estimation of structural networks, in a fully automatic manner. The estimated structural networks can be further analyzed based on potential associations between their degree of expression over subjects and different variables of interest (e.g. age, gender, site, etc.). Like in traditional SBM, this leads to fewer statistical tests. However, constrained SBM enables prior information to steer the decomposition and, thus, specific hypotheses to be investigated.

Both SBM and constrained SBM have been implemented on a common foundation where it assumes that data is locally available during running the experiments. But in neuroimaging studies, it is not always possible to pool all datasets in a centralized location. In addition, it can be time consuming and at the same time expensive as well to coordinate data collection and sharing across different sites. Even with proper permission controls and data anonymization, recent studies have found that there is potential risk of revealing the identity of subjects with rare diseases [12, 13]. The ensuing inability to pool datasets from their sites may eventually lead to poor overall computation performance since, sometimes, it may not be possible to detect disease effects from a single site’s low number of data samples. In recent years, there are lot of research ongoing to leverage decentralized computation for neuroimaging anlaysis [14, 15, 16, 17, 18]. Gazula et al. introduced a decentralized voxel based morphometry approach and applied it on 2000 structural magnetic resonance imaging datasets from 14 different sites [19], demonstrating the benefits of multiple variable analysis in a decentralized scenario. To overcome the difficulty in pooling datasets into one location, we introduce a novel method: decentralized constrained source-based morphometry (dcSBM). Our approach inherits multivariate properties from constrained SBM, but operates ICA across different sites in a distributed environment. At the end of decentralized evaluation, we aggregate loading parameters and sources from different sites and further investigate the obtained results with comparisons and ensuing statistical analysis.

In this work, at first, we describe the data acquisition and preprocessing steps. Following, we describe our dcSBM approach and evaluate its performance on real datasets. Here, we conducted comparative analysis between centralized and decentralized results on the same datasets. We investigated the reliability of our proposed model using a sizeable subset of the large sample UK Biobank prospective study. Then, we utilized the Function Biomedical Informatics Research Network (fBIRN) [20] and Center for Biomedical Research Excellence (COBRE) [21] structural MRI datasets to assess the clinical utility of this approach. The latter was done by assessing the significance and effect sizes of the association between the identified loading parameters of each source and different variables of interest. This work builds on our proposed method [22] and expands its application to clinical datasets, improving the statistical analysis of results and reliability assessments.

## 2. Methods

### 2.1. Data Acquisition and Preprocessing

We begin with a description of the three datasets utilized in this work.

#### 2.1.1. UK Biobank dataset

In UK Biobank dataset, sagittal orientation was used for the T1 structural MRI data. Following the previous measurements of population brain shape and size from [23], the front side of the brain is tilted down by 16° regarding anterior commissure - posterior commissure line. The T1-weighted structural image comprises the following parameters: field-of-view: 208 × 256 × 256 matrix, duration: 5 minutes, resolution: 1×1×1*mm*, 3D MPRAGE, sagittal, in-plane acceleration iPAT=2, prescan-normalize. The superior-inferior field-of-view is set to 256*mm* to capture the neck/mouth in a reasonable amount.

For UKBiobank dataset, we selected 3000 unaffected subjects for our experiments. The subject-level T1-weighted sMRI data were preprocessed using SPM12^1^ and a Gaussian kernel comprising FWHM = 10*mm* was used to smooth it. For the prior whole-brain references, thirty spatial references (*R*) were identified from separate group ICA results on two different sMRI datasets: 1) the genomics superstruct project (GSP), and 2) the human connectome project (HCP). Each dataset contains 3500 unaffected subjects [20].

#### 2.1.2. Function Biomedical Informatics Research Network (FBIRN) dataset

The Function Biomedical Informatics Research Network (FBIRN) datasets were collected from seven different sites. The subjects from six sites were collected on a 3T Siemens Tim Trio System while a 3T General Electric Discovery MR750 scanner was used for one of the sites. The T1-weighted acquisition parameters are described in [24].

The following subject inclusion criteria were applied to all subjects: head motion ≤ 3° and ≤ 3 mm. After applying these criteria, 165 healthy controls (HCs) (average age: 37.04 ± 10.86; range: 19-59 years; 46/119: female/male) and 152 patients with schizophrenia (SZs) (average age: 38.77 ± 11.63; range: 18-62 years; 34/118: female/male) were selected. Both HCs and SZs are matched by age, gender, and mean framewise displacement (FD) to avoid introducing confounding effects (age: p = 0.1758; gender: p = 0.3912; mean FD: p = 0.9657). It was ensured that 1) HCs do not have any past or current psychiatric illness based on SCID assessment or a first-degree relative with diagnosis of an Axis-I psychotic disorder, and 2) SZs were clinically stable at the time of scanning.

#### 2.1.3. Centers of Biomedical Research Excellence (COBRE) dataset

The MRI images were collected on a single 3T Siemens Trio scanner with a 12-channel radio frequency coil. T1-weighted high resolution structural images were acquired with a five-echo MPRAGE sequence with TE = 1.64, 3.5, 5.36, 7.22, 9.08 ms,

TR = 2.53 s, TI = 1.2 s, flip angle = 7°, number of excitations = 1, slice thickness = 1 mm, field of view = 256 mm, resolution = 256 × 256.

Like fBIRN, we applied similar subject inclusion criteria to the COBRE dataset, resulting in 82 HCs (average age: 38.09 ± 11.67; range: 18 to 65 years; 22*/*60: female/male) and 66 SZs (average age: 37.79 ± 14.45; range: 19 to 65 years; 9*/*57: female/male). HCs and SZs are also matched by age, gender and mean FD (age: *p* = 0.8874; gender: *p* = 0.0794; mean FD: *p* = 0.6475).

For fBIRN and COBRE datasets, we preprocessed the sMRI data using SPM12. We segmented the structural images into 6 tissue compartments, including gray matter, white matter, and CSF, using modulated normalization, resulting in gray matter volume (GMV) features. These were then smoothed by a Gaussian kernel comprising the full width at half maximum (FWHM) = 6 mm.

### 2.2. Spatially Constrained ICA

ICA is a widely used approach in biomedical domain [25, 26, 27, 28]. In structural MRI data, at first, ICA is imposed on subject volume matrix *X*, and decompose into a mixing matrix *A*, and a source matrix *S*. The mixing matrix represents the relationship between subjects and sources, and source matrix preserves the relationship between sources and voxels of brain. The ICA decomposition applies special filtering to handle noise and provides maximally spatially independent sources which pertains significant inter-subject covariation [29].

Constrained ICA is an extension of ICA which utilizes prior information during the processs of decomposition and extracts targeted sources. For the prior information, at first a reference *R* which contains desired sources is chosen. An augmented Lagrange multiplier is utilized for the identification of desired sources, the closest to the reference parameter is enabled and finally maximally accurate *S* is selected [30]. A closeness measure *ε*(*S, R*) between an obtained signal *S* and the reference signal *R* is defined to restrict the learning process. So, only single weight will be found to give the source *S*, which is closest to the reference signal *R*. This constrained ICA framework is formulated as follows [31]:

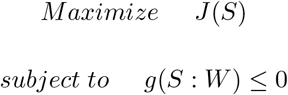

*Maximize J*(*S*) *subject to g*(*S* : *W*) ≤ 0 Here *J*(*S*) denotes the contrast function of a standard ICA algorithm, *g*(*S* : *W* = *ε*(*S, R*) − *ξ* ≤ 0.

### 2.3. Decentralized Source Based Morphometry

To utilized the multivariate constrained SBM approach on a dataset that is distributed to different local sites, we introduce decentralized constrained SBM (dcSBM). In the centralized case, we have a dataset *X* = [*x*_1_ …, *x*_*N*_]^T^, where *x*_*i*_ ∈ R^*V*^ is the *V* dimensional feature vectors of *i*-th subject. Now we apply constrained SBM on this dataset and extract gray matter differences. In the decentralized environment, there are total *L* sites, and each site *l* has dataset *D*_*l*_ = {(*x*_*i*_, *z*_*i*_) : *i* ∈ {1, 2, …, *N*_*l*_}} from *N*_*l*_ subjects, where *z*_*i*_ ∈ R expresses the predictor variables age, gender, diagnosis, and site from subject *i*. Here, each local site *l* operates constrained ICA separately on their local data *X* and stores mixing (*A*_*l*_), and source matrices (*S*_*l*_). In mixing matrix, each column is called loading parameters. Before running local ICA, each site utilizes the same reference map *R* which contains the prior information about the desired sources. The constrained ICA was applied on the data matrix *X*, guided by a fast fixed-point algorithm. The is included as part of the group ICA toolbox GIFT^2^. Guided by the reference vector *R*, the local source matrix *S* and mixing matrix *A* are obtained from the data matrix *X*.

In mixing matrix *A*, the scores in each columns represent the contribution of each component to all subjects. Meanwhile, the scores within row of source matrix *S* represents statistically independent spatial configurations that specify the areas of coherent variability among subjects. We operated constrained ICA to each site, and finally, a master node concatenated the mixing matrix from each site by the decentralized aggregation process. For the quality assessment of our approach, the source matrices were also aggregated from each site. The flow diagram of our approach is shown in Fig 1.

**Figure 1:**
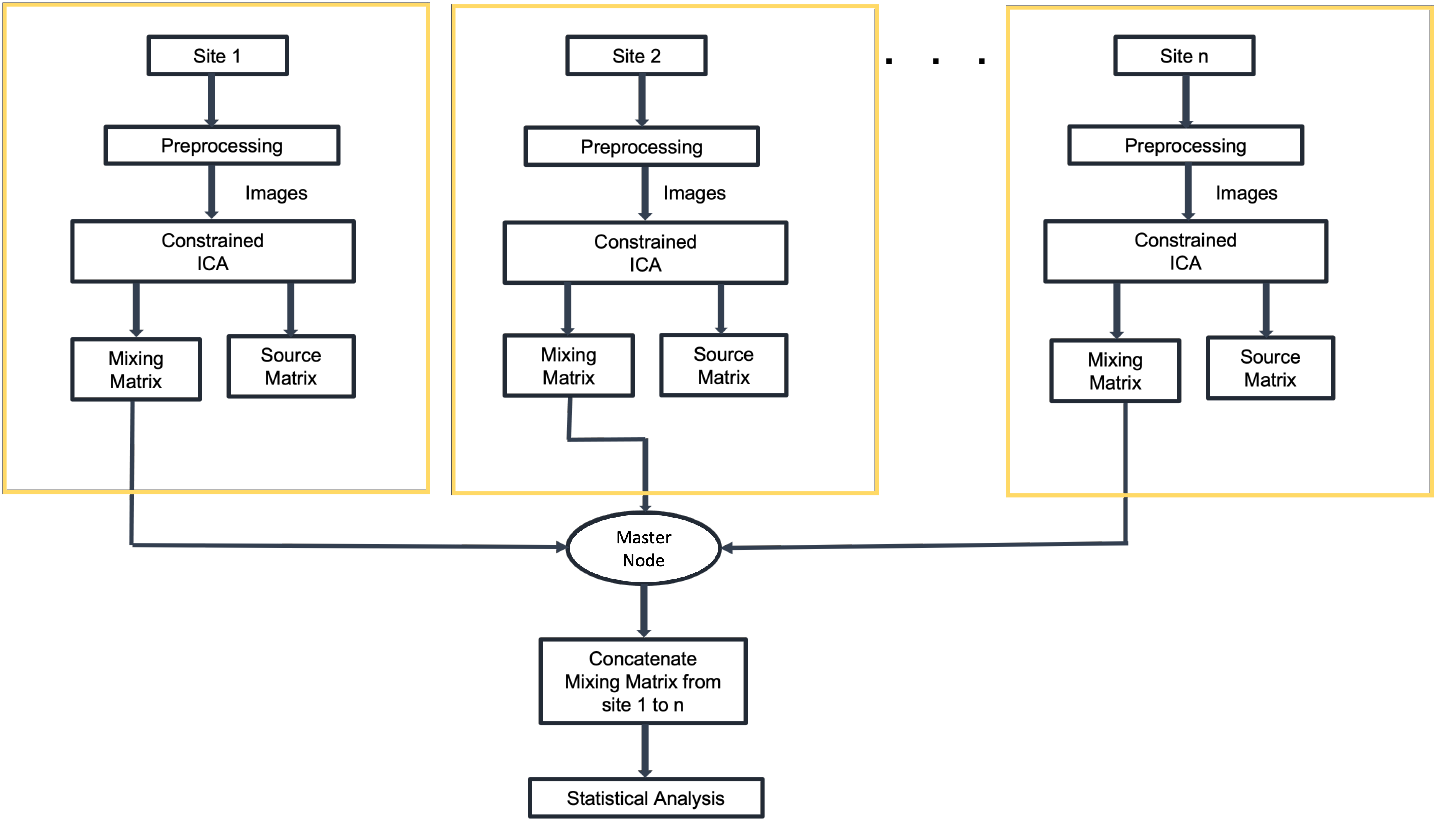
Flow diagram of decentralized constrained SBM

To perform the statistical analysis, we also operate constrained ICA in a centralized environment. In centralized experiment, we pool data from all sites together and run ICA on this. We extracted mixing, and source matrix; and finally compared with the decentralized results.

### 2.4. Statistical Analysis

#### 2.4.1. Pairwise Correlation

In this statistical analysis, we used fBIRN and COBRE dataset. As we acquired fBIRN dataset from seven different sites, we consider each of them as individual local site during running dcSBM. We collected COBRE dataset from a single site, and consider this as a single site during the computation. Finally, there are eight different sites participated in this analysis. After running constrained ICA to each local site separately, we concatenated the mixing matrix, and aggregated the source matrix from each site by a common aggregator. To analyze the decentralize results, we also pooled all fBIRN and COBRE datasets together and run constrained ICA centrally. Then, we evaluated the correlation between *i*^*th*^ column of centralized loading and *j*^*th*^ column of decentralized loading parameters (where *i* == *j*). Finally, We also computed the correlations between each column of centralized and decentralized source metrices.

#### 2.4.2. Linear Regression

We applied linear regression on decentralized SBM model. Like Pairwise correlation, we use fBIRN and COBRE datasets here. Our predictor matrix *Z* has the dimensions with 465 × 10, where the variables are age, gender, diagnosis, site, and interactions between gender and diagnosis, gender and age, diagnosis and age, age and site, diagnosis and site, and gender and site. Here we used loading parameters as our response variable. We utilized the same mixing matrix that we used in computing pairwise correlation(Section 2.4.1). Finally, we fit our data matrix *Z* to a linear regression model. As we have 30 loading parameters columns(dependent variables), we run our linear regression model 30 times and store the results. For the comparative analysis, we also applied linear regression on the centralized SBM model.

We also computed the effect of independent variables to dependent variables. In our analysis, we use loading parameters as the dependent variables and consider age, gender, diagnosis, sites and possible interactions between them as the independent variables. We used Anova Type ⨿ method to evaluate the effect size.

#### 2.4.3. Model reliability

For examining the reliability of our proposed method, here we design two different experiments using the UK Biobank dataset. For the first experiment, we calculate the correlation, Mean Square Error (MSE), Max Absolute Error (MaxAE), and Median Absolute Error (MedianAE) among the correlation matrices of loadings, and sources separately. For the second experiment, we compute correlation, and Mean Square Error (MSE) between the correlation matrices of loadings, and sources separately.

### I Random shuffle

In this experiment, we artificially created three different sites and 3000 UK Biobank subjects were used to perform this experiment. In every experiment, we shuffle the subjects and assigned 1000 distinct subjects to each site. We repeated this experiment ten times and collected the results for the statistical analysis.

### II Increase sites with constant subjects

In this experiment, we ran five different experiments. Every time we increase the number of local site by one and assign constant 500 distinct subjects to this site. The five different experiments consist of 2, 3, 4, 5, and 6 different sites respectively. In each experiment, we also pooled subjects from the local site and run the analysis centrally to compare with the decentralized results.

## 3. Results

In dcSBM, we extracted thirty independent components by applying spatially constrained ICA to each site separately. We also pooled all dataset together from each site and run spatially constrained ICA on this in a centralized environment. Finally, we compare our decentralized results with the centralized estimates to validate our model. Thirty loading parameter columns (one for each maximally independent component) are included in the mixing matrix. We take centralized and decentralized loading parameters and computed the correlations between them. The experimental results are presented in Fig 2. Panel (A) represents the correlation among the centralized loading parameters only, and panel (B) stands for the correlation among the centralized and decentralized loading. In centralized vs decentralized correlation plot, we find the high correlation score across the diagonal which implies that the loading parameter *i*, acquired from centralized SBM and loading parameter *j* comes from decentralized SBM (where *i* == *j*) are highly correlated. The correlations between centralized and decetralized source matrix are presented in Fig 2(C), and (D). In Fig 2, panel (C) presents the correlation among the centralized sources, and panel (D) is the correlation plot between the centralized and decentralized source matrix. We also observe the same correlation pattern where we obtain high correlation across the diagonal.

**Figure 2:**
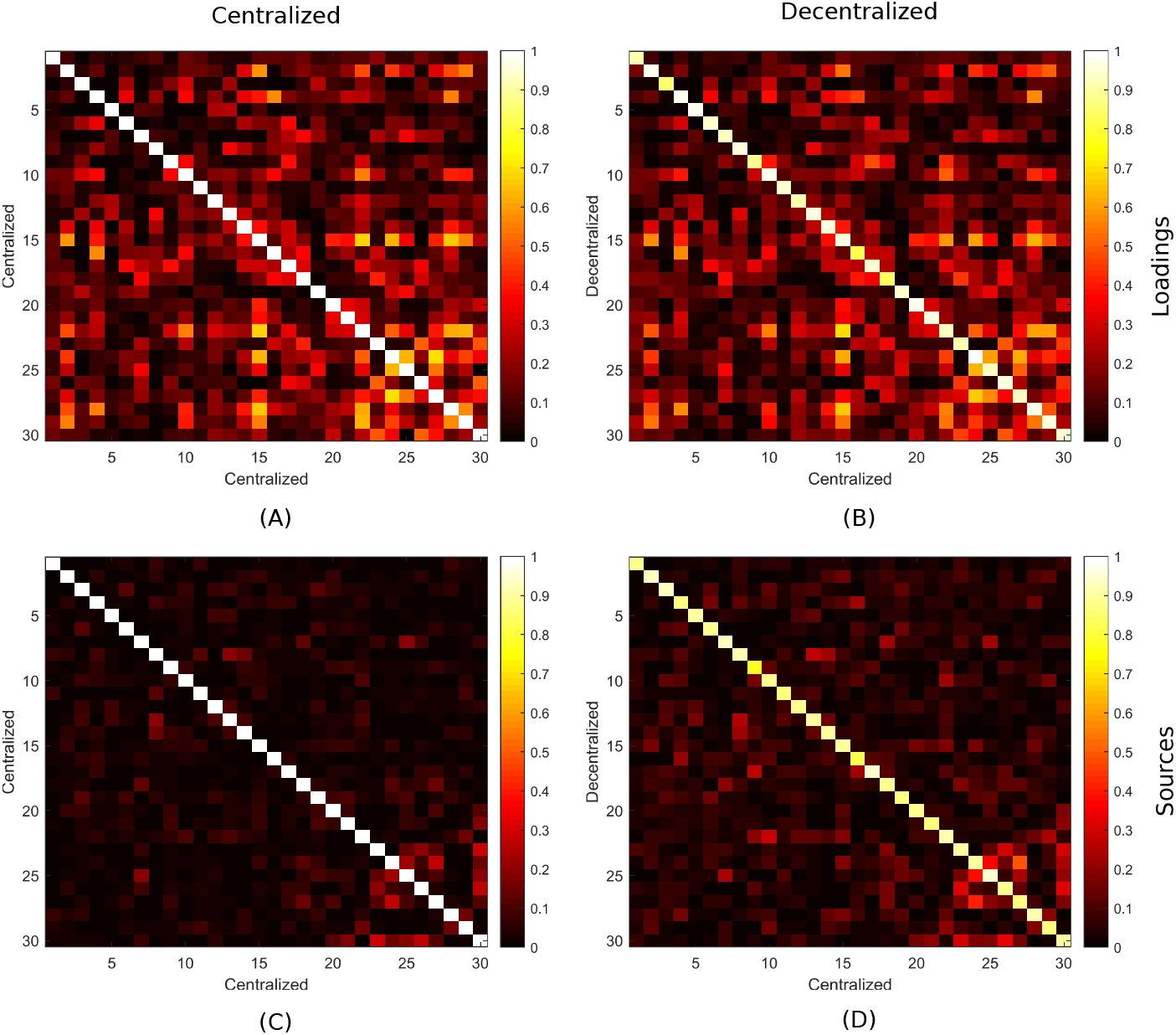
Correlation plots of loading parameters and sources. Panels (A), and (B) represent the correlation between centralized versus centralized, and centralized versus decentralized loading parameters respectively. Panels (C), and (D) represent the correlation between centralized versus centralized and centralized versus decentralized sources respectively. Panels (A), and (C) are the expected similarity structures for the centralized loadings and sources respectively. Panels (B) and (D) are the recovered similarity structures among the centralized and decentralized loadings and sources respectively. The diagonal exhibits the near-1 correlations.

In the following regression analysis, we consider loading parameters as our dependent variable and use age, gender, diagnosis, site, and possible interactions between them as independent variables (Section 2.4.2). From centralized SBM and our dcSBM results, we observed a large effect size of diagnosis for several loading parameters. We observed the largest effect sizes for loading parameters 4 and 28. In Fig 3, we present cloud plots of loadings 4 and 28 using the adjusted loading parameter values obtained from the linear regression model. In each panel, there are two cloud plots. The top one is the cloud plot of adjusted loading parameters for the schizophrenia patients and bottom one is for the healthy controls. In Fig 3, panels (A), and (B) present the cloud plot of adjusted values of loading parameter 4 for the centralized and decentralized analysis, respectively. Similarly, panels (C) and (D) are the cloud plots of loading 28 for the centralized and decentralized analysis, respectively. From the figure, we observe that the centralized and decentralized loading scores are very similar in shape, but differ in scale.

**Figure 3:**
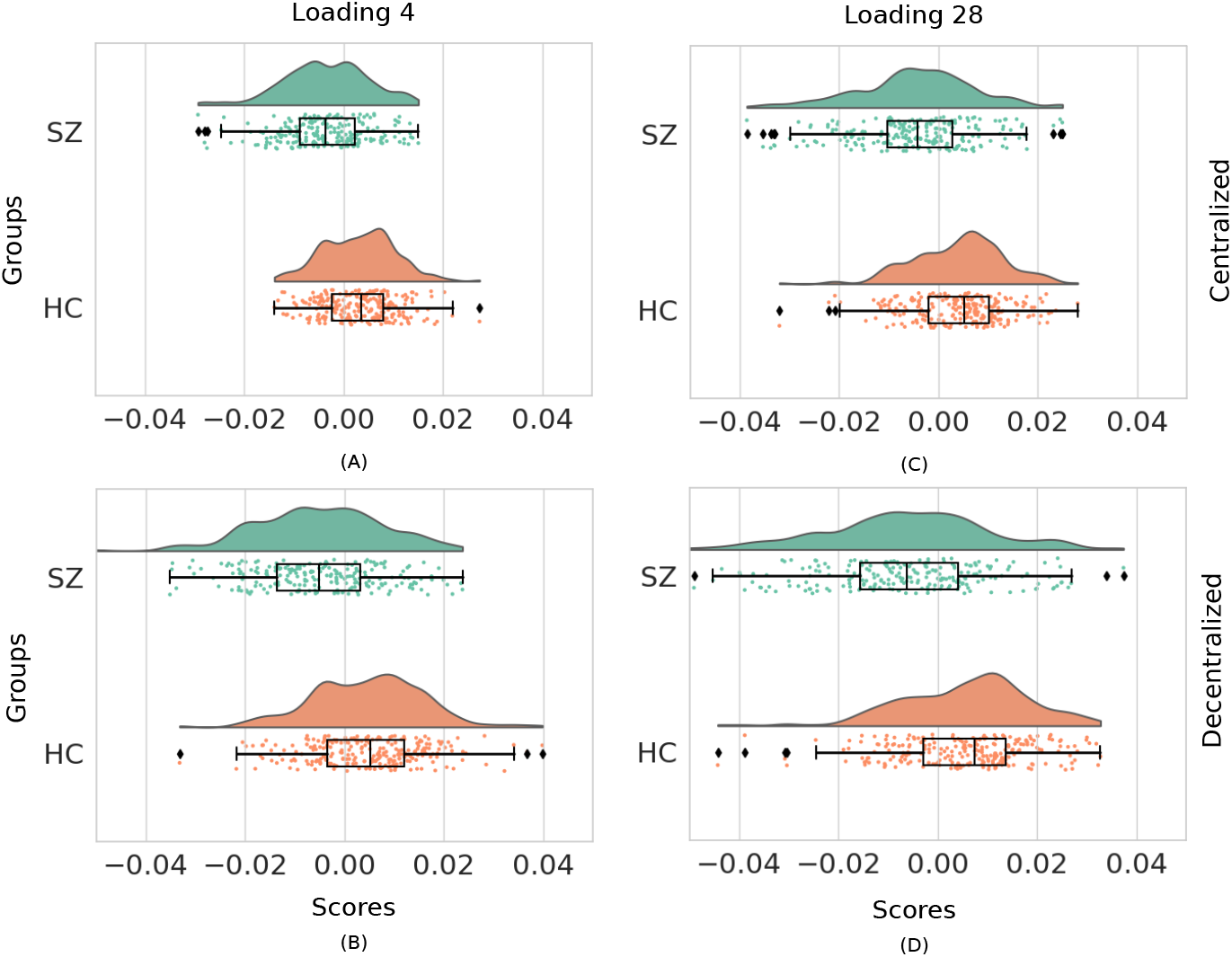
Cloud plot of adjusted loading parameters 4 and 28. Panels (A) and (B) represents the cloud plot of adjusted loading parameters 4 for the centralized and decentralized analysis respectively. Panels (C) and (D) presents the plot of adjusted loading parameters 28 for the centralized and decentralized analysis respectively. In each panel, upper plot(Green) and lower(Brown) plots are the clouds of loading score for the patients and healthy controls respectively.

From our regression model, we computed the effect size for independent variables age, gender and diagnosis for both centralized and decentralized analysis. We present our results in Fig 4. In the figure, panels (A), (B), and (C) are the scatter plots of the centralized vs decentralized effect sizes for the variables age, gender, and diagnosis respectively. In each panel, we observe points largely follow the identity line. These plots demonstrate that the obtained centralized and decentralized effect sizes are very similar. Based on the effect sizes of diagnosis for the centralized case being *ϵ*^2^ *>* 0.02, we identified seven independent sources. The spatial maps of these sources are presented in Fig 5. Notice that although the effect sizes in the decentralized case tend to be slightly inflated with respect to the centralized case, only a couple of false positives occur for diagnosis (our effect of interest). We also computed the P value for variables age, gender, and diagnosis obtained from our linear regression model and finally compared decentralized results with the P values from the centralized analysis. We present the experimental results in Fig 6. In the Figure, panels (A), (B), and (C) demonstrates the scatter plots of centralized vs decentralized P values for the variables age, gender, and diagnosis respectively. Here we find very similar *P* values for centralized and decentralized analysis and in close correspondence with the patters observed for effect sizes.

**Figure 4:**
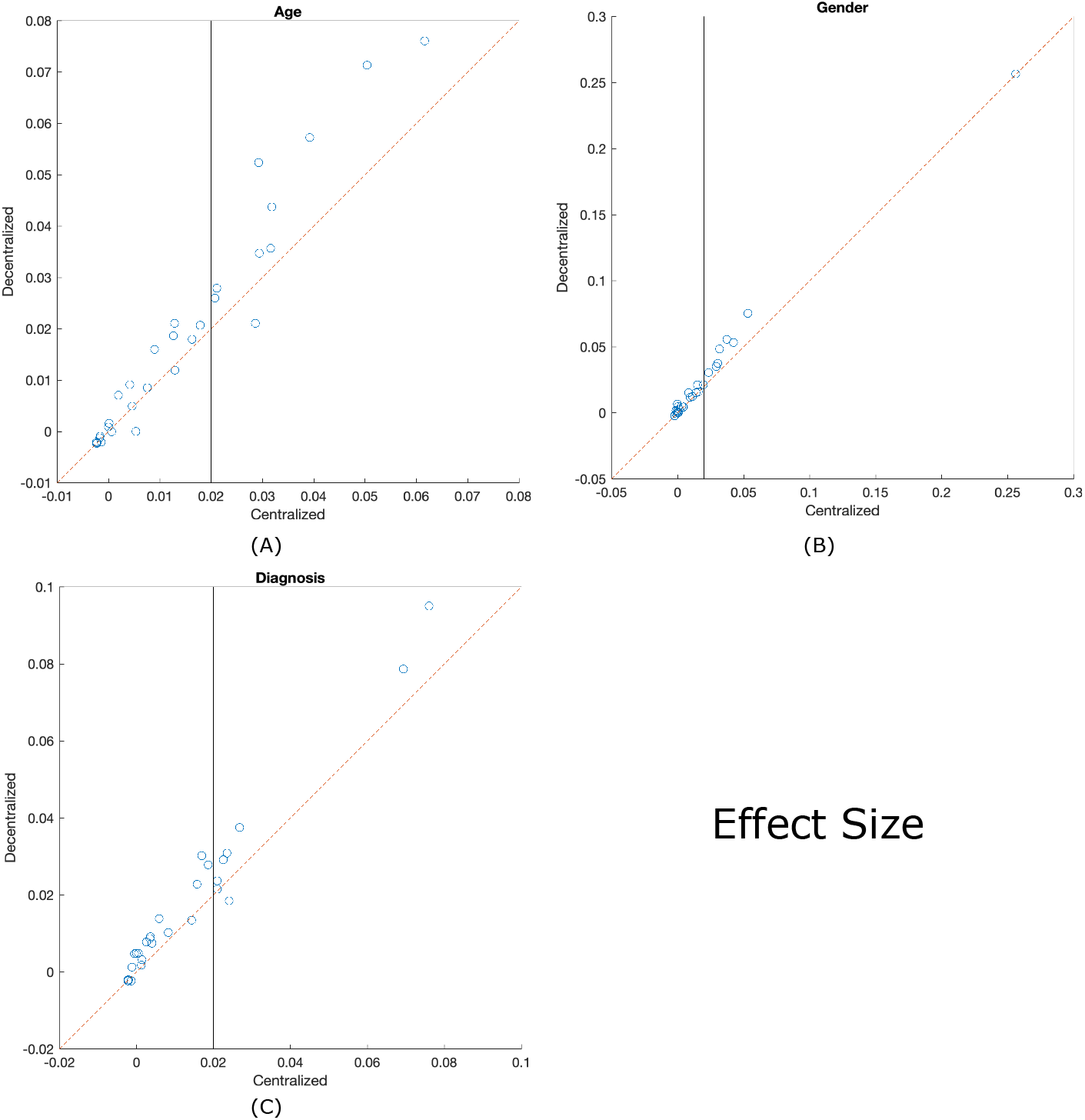
Scatter plot between the centralized and decentralized effect sizes. Panels (A), (B), and (C) correspond to variables age, gender, and diagnosis, respectively. Scatter plots of centralized and decentralized effect sizes exhibit high consistency.

**Figure 5:**
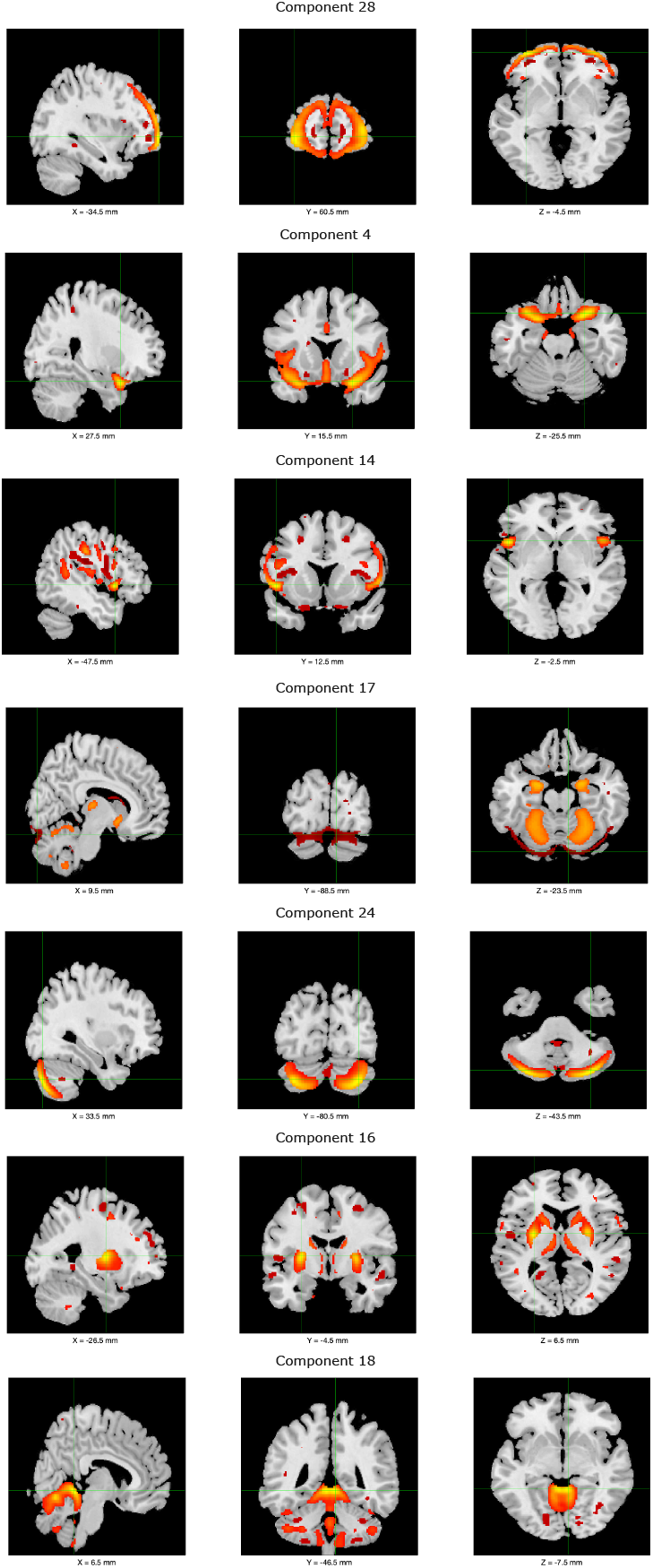
Visual summary of sources with large effect sizes

**Figure 6:**
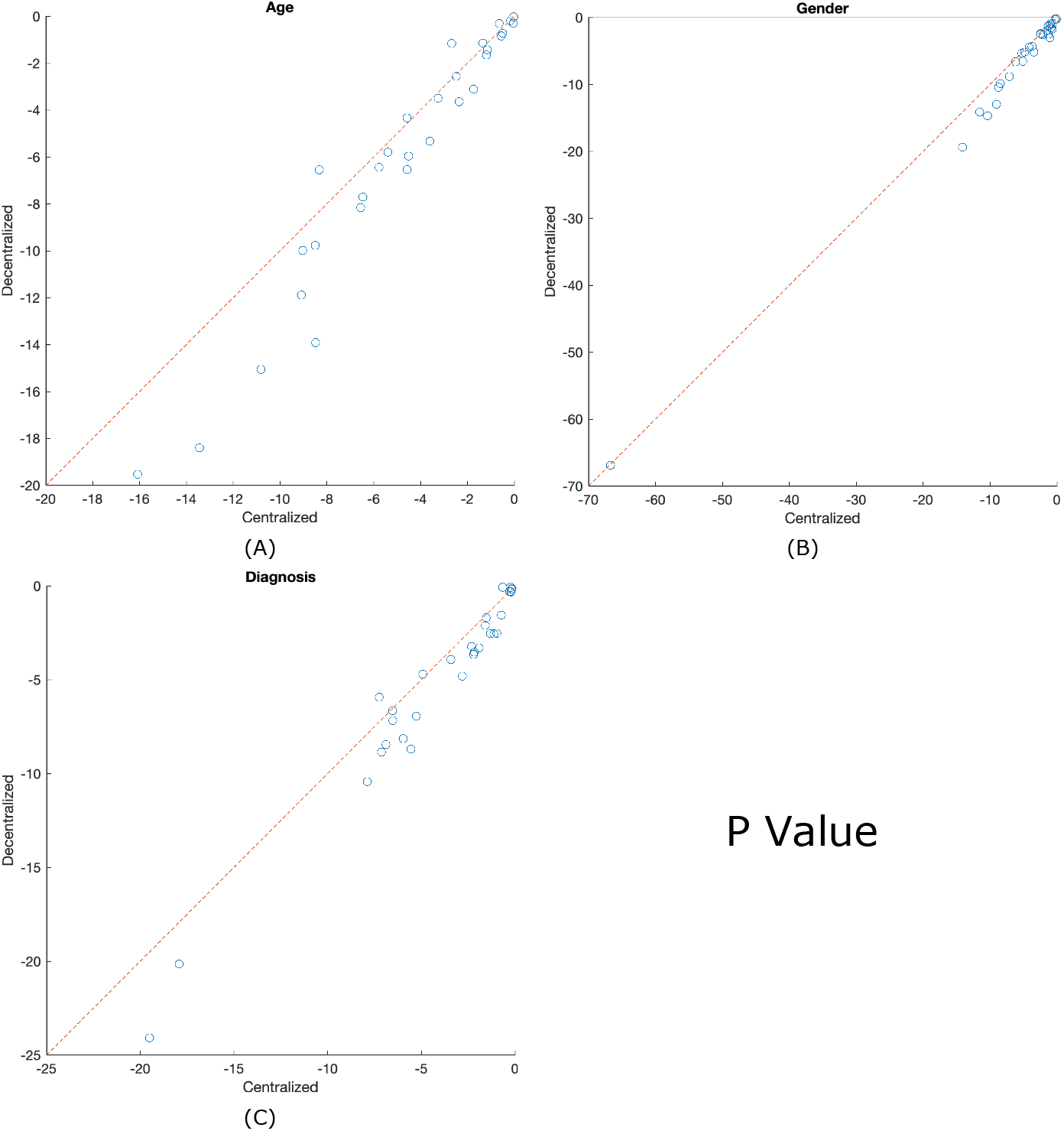
Scatter plot of centralized versus decentralized P values. The values are presented in log-log scale. Panels (A), (B), and (C) correspond to variables age, gender, and diagnosis, respectively. The similar *P* values for centralized and decentralized analysis indicate high consistency.

To examine the reliability of our dcSBM approach, we utilized the UK Biobank dataset. We present the experimental results in Fig 7 (Section 2.4.3: random shuffle). Fig 7(A) presents the similarity among the centralized and decentralized loading parameters. The performance measures correlation, mean square error (MSE), max absolute error (MaxAE), and median absolute error (MedianAE) are among the correlation matrices shown in Fig. 2(A) and Fig. 2(B) obtained for the UKBiobank dataset, for each of the 10 shuffled runs. Here each MSE, MaxAE, and MedianAE are normalized by the mean-squared, max-abs, and median-abs of centralized correlations matrices, respectively. Fig 7(B) represents the similarly between the centralized source correlation matrix and centralized versus decentralized correlation matrix. In the Figure, Each boxplot consists of 10 points from different 10 shuffled runs, and we find all points are very similar, which eventually implies high reliability.

**Figure 7:**
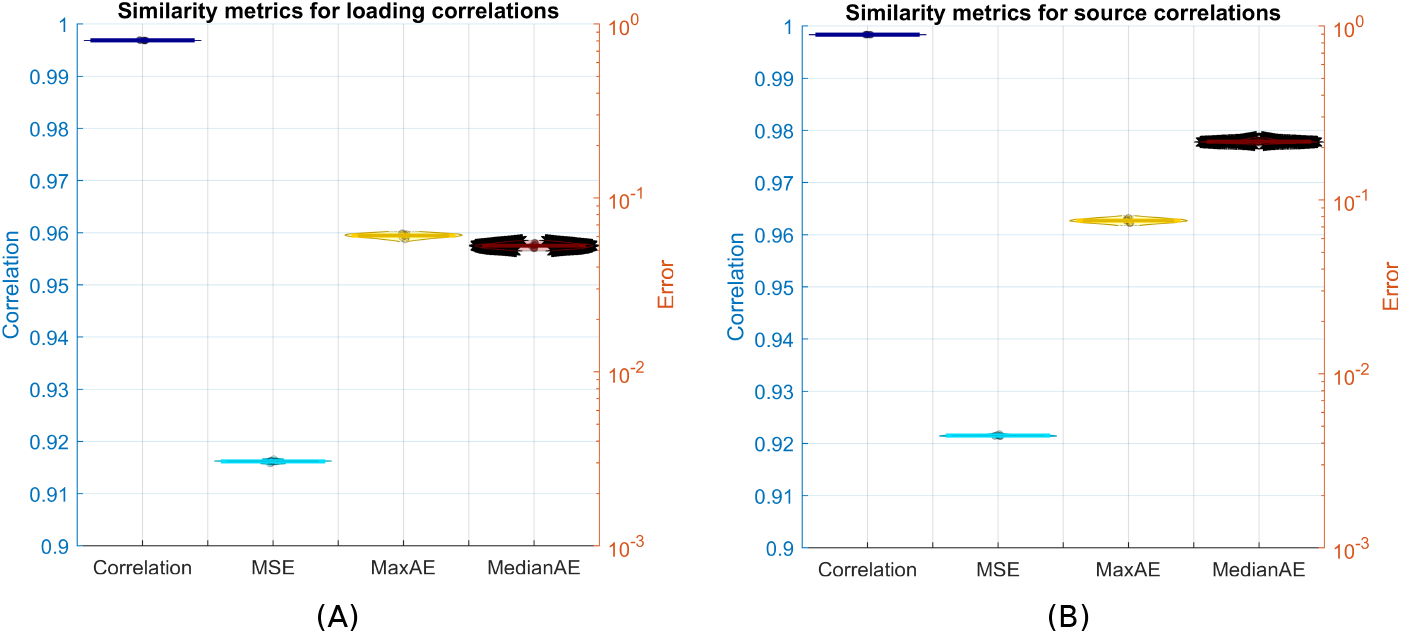
Similarity metrics of loading parameters and sources. Panels (A), and (B) present the similarity metrics of loading parameters, and sources respectively. The left Yaxis represent the correlation measure and right Yaxis present mean square error, max absolute error, and median absolute error.

Fig 8 represents the results of Section 2.4.3: increasing number of sites, each with the same number of subjects. After running the experiment for each site, we collect the correlation matrices (as in Fig. 2(A) and Fig. 2(B)) and report the correlation and MSE between them. We also compute these metrics for the source correlation matrices. Results are presented in Fig 8 (A) and (B), respectively. In both plots, we observe very high correlation between the centralized and decentralized results, with the correlations decreasing and MSE increasing slightly as the total number of sites participating in the analysis increases.

**Figure 8:**
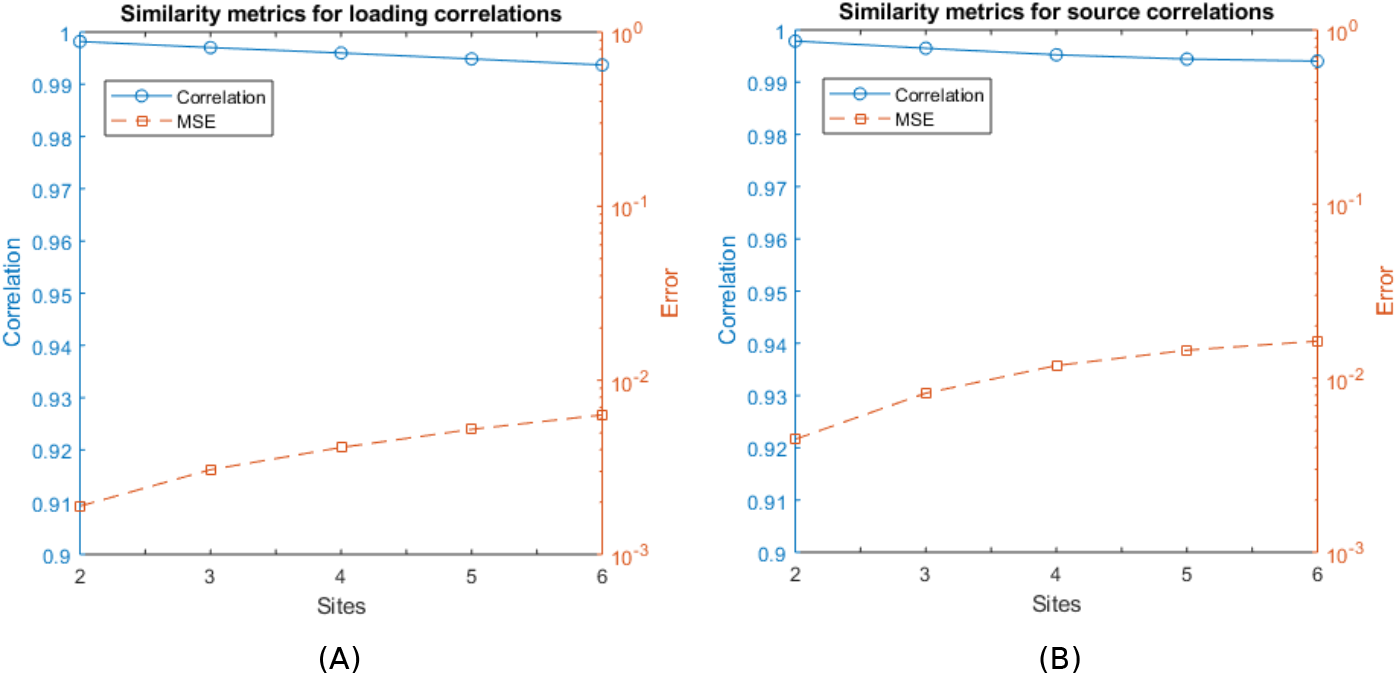
Similarity metrics for loading parameters and sources. Panels (A), and (B) represent the similarity metrics of loading parameters, and sources respectively for six different sites. The left Yaxis, and right Yaxis are utilized for correlation, and mean square error (MSE) respectively.

## 4. Discussion

Interest in utilizing sMRI data to analyze brain morphometry continues to grow. SMRI features have successfully been applied to examine the structure and function-alterations in different domains [32, 33], patient classifications [34, 35, 36], and in a multimodal study [37, 38]. In a scenario where data is only locally accessible and cannot be pooled in a central location, many established algorithms, like constrained SBM, can not operate. To overcome the limitation of constrained SBM, we have introduced our decentralized constrained SBM algorithm to operate on decentralized environments.

In our analysis, thirty independent components were extracted from both centralized and decentralized SBM. To analyze the pairwise correlation, we computed the correlation among each column of the centralized and decentralized loading parameters. For comparative analysis, we also evaluated the correlations among the centralized and decentralized source matrices. The results are presented in Fig 2. From our statistical analysis, we observed that the centralized and decentralized result estimates are very similar. That’s why in the correlation metric, the diagonal exhibits highest correlations. We also observed that if the centralized estimates extracts any structural patterns, these patterns are also reflected to our decentralized results.

Recently, many studies have been conducted to analyze the effects of age, gender, and diagnosis [24, 39, 40, 41, 42, 43, 44, 45] on gray matter. In our regression analysis, we consider age, gender, diagnosis, site, and the possible interactions between them as the predictors. The loading parameters recovered from local constrained ICAs are considered as the dependent variables. We computed the *p* value for the variables age, gender and diagnosis for both centralized and decentralized SBM. The results are presented in Fig 6. We computed the minimum, maximum, and standard deviation from the obtained *p* values. We summarize the centralized vs decentralized *p* values in Table. 1. In the table, c−min, d−min, c−max, d−max, c−std, and d−std represent the centralized minimum, decentralized minimum, centralized maximum, decentralized maximum, centralized standard deviation, and decentralized standard deviation respectively. The results are mostly aligned across the diagonal. We also computed the effect size of variables age, gender, and diagnosis to loading parameters. The results are presented in Table. 2. Here the results from both centralized SBM and our dcSBM exhibit similar patterns where the plots align across the diagonal. All-in-all, these experimental results establish the hypothesis that our dcSBM produces very close results compared to centralized SBM.

**Table 1:** Centralized vs decentralized p values

| variables | c-min | d-min | c-max | d-max | c-std | d-std |
| --- | --- | --- | --- | --- | --- | --- |
| Age | 0.0000 | 0.0000 | 0.9846 | 0.9685 | 0.3084 | 0.2908 |
| Gender | 0.0000 | 0.0000 | 0.9505 | 0.8132 | 0.2790 | 0.2426 |
| Diagnosis | 0.0000 | 0.0000 | 0.8507 | 0.9377 | 0.2997 | 0.3329 |

**Table 2:** Centralized vs decentralized effect sizes

| variables | c-min | d-min | c-max | d-max | c-std | d-std |
| --- | --- | --- | --- | --- | --- | --- |
| Age | -0.0023 | -0.0023 | 0.0616 | 0.0761 | 0.0168 | 0.0223 |
| Gender | -0.0023 | -0.0022 | 0.2562 | 0.2567 | 0.0471 | 0.0483 |
| Diagnosis | -0.0022 | -0.0023 | 0.0760 | 0.0950 | 0.0191 | 0.0225 |

We conducted these experiments on real data (seven sites for fBIRN and single site for COBRE). We also considered the site effect in our regression analysis, including its interaction with age, gender, and diagnosis. Based on the effect sizes from diagnosis, we identified seven components whose effect sizes are significant. Component 4 covers the insula, temporal pole, frontal orbital cortex, anterior cingulate, and parahippocampal domains; Component 28 covers the regions of anterior and dorsolateral prefrontal cortex; Components 14 covers Insula and parietal areas; component 17 covers cerebellum and subcortical areas; component 24 covers additional regions of cerebellum; component 16 covers putamen and caudate nucleus (subcortical), and component 18 covers vermis and declive of the cerebellum. Previous studies also reported similar component regions [24, 46, 47, 48, 49], which further supports the validity of our findings. The references used in the patient dataset can be organized into eight different domains: five components in Visual (VS), nine in Cerebellum (CB), three in Frontal (FN), two in Default-Mode (DMN), two in Sub-cortical (SC), two in sensorimotor (SM), two in Insula (IS), and one in Hippocampus (HIP) domain. The spatial maps of eight different brain regions from our analysis are presented in Fig 9.

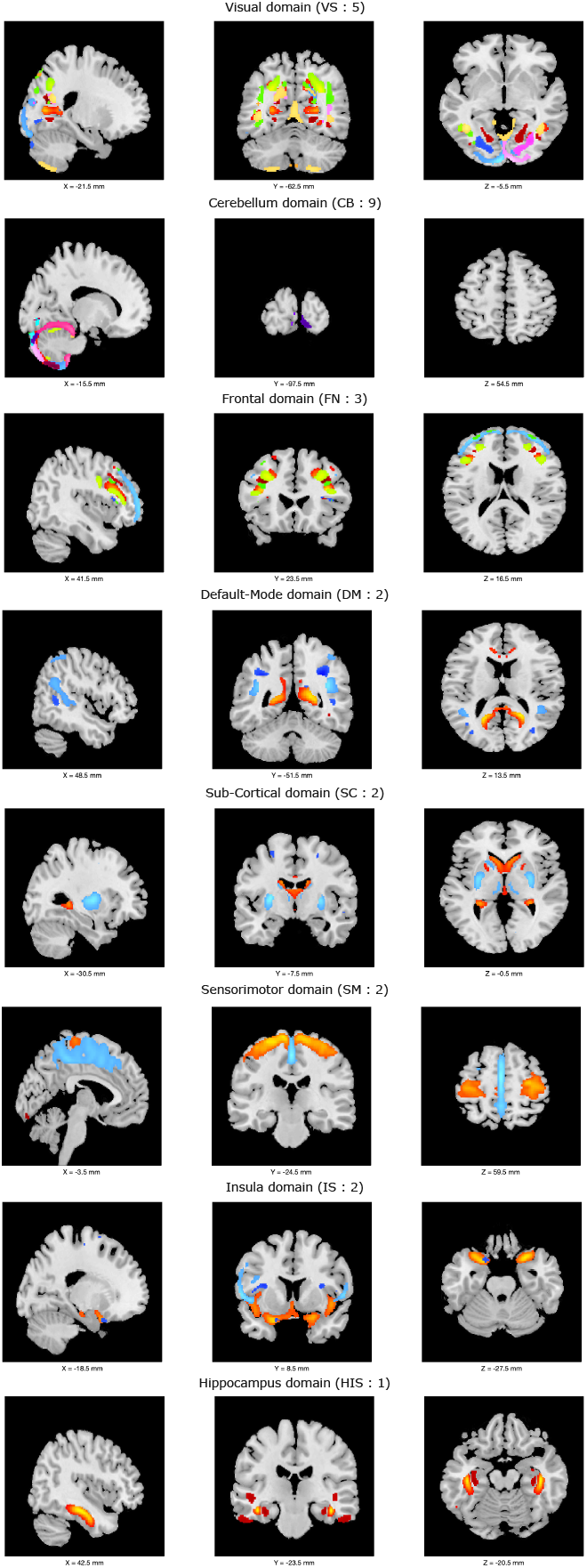
Visual summary of components in eight different brain regions: five components in Visual (VS), nine in Cerebellum (CB), three in Frontal (FN), two in Default-Mode (DMN), two in Sub-cortical (SC), two in sensorimotor (SM), two in Insula (IS), and one in Hippocampus (HIP) domain

Lastly, we acknowledge that the estimated loading parameters produced with dcSBM have larger values than the centralized counterparts. This is possibly due to the know scaling ambiguity in ICA. While this could pose issues such as inflation of site effects in our proposed decentralized setting, our statistical analysis modeled such effects in a way that mitigates these concerns (i.e., using linear an interaction terms with site).

### 4.1. Decentralized regression without looking loading parameters

In our approach, a master node accumulates the mixing and source matrices from each site to operate the statistical analysis at a central site. Our aggregate statistical operation may raise privacy concerns. However, note that our main goal was to establish the hypothesis that the decentralized constrained SBM can very closely replicate the centralized estimates. In our analysis, we concatenated loading parameters from each site in an aggregator and ran regression analysis centrally. But, it is possible to conduct the linear regression analysis in a totally distributed manner. For decentralized regression, we refer the reader to [50]. In summary, we can write the least squares solution for the regression parameter 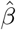as:

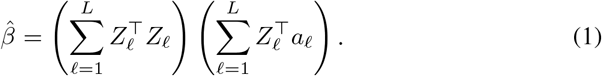

In (Equation 1), *a*_*l*_ is the response variable we would like to predict, here the loading parameters learned locally in site *l* based on the linear model *a* ≈ *Zβ*, where *Z* is the matrix containing predictors in each column, and *β* is the regression parameter. In a decentralized environment with *L* sites, *Z* can be represented by 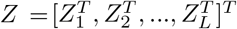, and *a* can be defined by 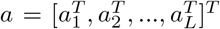 . In this approach, *a*_*l*_ and *Z*_*l*_ do not have to be transmitted/aggregated, improving the privacy. Only the terms 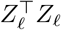 and 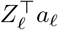 are transmitted/aggregated. This approach is capable of generating identical results as our simple aggregator and has been integrated to the Collaborative Informatics and Neuroimaging Suite Toolkit for Anonymous Computation (COINSTAC^3^) [51, 52]. COINSTAC is a dynamic and decentralized software platform for decentralized computations that provide privacy-preserving features. Lastly, it is worth to mention that in our approach, the source matrices were aggregated from each site only for the purpose of quality control assessment of our proposed approach. It is recommended that each site retain their source estimates (*S*_*l*_) to minimize risks of privacy leaks (note that, by definition, the local data *X*_*l*_ can be reconstructed from *A*_*l*_ and *S*_*l*_).

## 5. Conclusion

In this work, we have introduced a novel approach to operate the constrained SBM in decentralized manner. We have utilized two different real multi-site datasets for our statistical analysis. In our experiment, we considered age, gender, diagnosis, and site effects and evaluated the similarity between centralized and decentralized constrained SBM. We also investigated the reliability of our algorithm using the UK-Biobank dataset. In all experiments, our decentralized approach generates very close results compared to centralized estimates. Based on the group differences, we identified two sources, one covering the insula, temporal pole, frontal orbital cortex, anterior cingulate, and parahippocampal domains, and another covering the anterior and dorsolateral prefrontal cortices. For future work, we will deploy a decentralized regression approach to conduct our experiment without results aggregation. We will also integrate our dcSBM approach within the COINSTAC application to make it openly available to the community. Finally, our algorithm gives the benefit to the private local site data which needs to stay locally while still leveraging its utility via a decentralized computation.

## Acknowledgement

This work was funded by the NIH (R01DA040487) and NSF (2112455). This research has been conducted using the UK Biobank Resource under Application Number

## Footnotes

1 https://www.fil.ion.ucl.ac.uk/spm/software/spm12/

2 http://trendscenter.org/software/gift

3 https://coinstac.org/

